# Effects of variations in adaptation potential on invasion speeds and species ranges

**DOI:** 10.1101/529735

**Authors:** José Méndez-Vera, Gaël Raoul, François Massol, Nicolas Loeuille

## Abstract

Confronted with global changes and their potential impacts on biodiversity, an important question is to understand the ecological and evolutionary determinants of species geographical distributions. In order to understand how adaptation in heterogeneous environments constrains such distributions, we analyze how the potential of adaptation along an environmental cline affects the geographical distribution and propagation dynamics (invasion or extinction) of a single species. We re-analyse a model initially proposed by Kirkpatrick and Barton using propagation speed to assess whether species distribution is spatially limited or not.

We found that for big adaptation potentials, the species invades space following Fisher’s model, whereas for small adaptation potentials the propagation depends on the evolutionary challenge to overcome. We have explicit approximations for the propagation speeds in both cases. We discuss the utility of these propagation speeds as an eco-evolutionary index based on empirical studies.

## 1. Introduction

In response to current climate changes, many species have been observed to shift their geographic distribution (Parmesan and Yohe (2003)). Such changes in the spatial distribution of species may largely alter their co-occurence, thereby affecting the structure of ecological networks (Tylianakis et al. (2008)), the functioning of ecosystems and the services they provide. To better understand such consequences, we urgently need to predict how species establish themselves along environmental gradients, but also to understand the mechanisms determining species distributions, so that we can forecast their future changes and thus adapt conservation policies.

To tackle this question, the most common approach relies on the development of niche-based species distribution models (SDMs), which provide predictions of species distributions based on presence/absence data and their association with a given set of environmental variables (refer to Guisan and Zimmermann (2000) for an introduction to SDMs or to Guisan and Thuiller (2005) for a more recent review; see also Thuiller et al. (2003) for a comparison of the performance of some SDMs). SDMs usually assume niche conservatism and range equilibrium, thus failing to include local adaptation. On some occasions, niche models alone may fail to describe the observed distribution of a species, especially in out-of-equilibrium cases such as species invasions. For instance, Broennimann et al. (2007) document a case study in which an invasive species has a different niche in its invasion range, although in this case data does not allow to determine if differences are adaptive (due to a shift in fundamental niche) or ecological (due to another possible realized niche taking place). Understanding such aspects would require the development of models which would simultaneously consider the niche model and a mechanistic approach of eco-evolutionary dynamics (e.g., Bush et al. (2016)).

Although adaptation to local conditions should help a population expand its range, boundary populations may be constrained in their adaptation due to the negative effect of gene flows from more central populations, i.e. genetic swamping. For example, Sanford et al. (2006) and Dawson et al. (2010) observed high migration load in boundary populations of a fiddler crab (*Uca pugnax*) and a volcano barnacle species (*Tetraclita rubescens*), respectively, while showing that individuals from the range limit are able to produce offsprings that would survive past the limit. Adaptation may take place on relatively short timescales: Balanya (2006) has shown that *Drosophila subobscura* at the leading edge of an ongoing invasion are able to adapt to local conditions while establishing a cline of genetic characteristics linked to temperature adaptation, following climatic gradients. Rapid adaptation and genetic swamping are quite general phenomena not restricted to species with short generation time. High gene flow has for instance been suggested to occur in many tree species Kremer et al. (2012), with potentially important effects on genetic variance at edge populations. Such evolutionary constraints may play a critical role in the persistence of tree species and in the variations of their geographic distributions, affecting the future of forest ecosystems under climate change scenarios. These studies underline the crucial need of including local adaptation when studying species distributions, and even more so when the aim is to understand and forecast future distributions under global change.

One monospecific spatially structured model accounting for both local adaptation and migration was presented by Kirkpatrick and Barton (1997). This model explains limited range distribution as an equilibrium between migration and genetic load from maladapted populations. Since there are no known explicit solutions, this model needs to be applied through numerical simulations.

In the present work, we take another look at the model by Kirkpatrick and Barton (1997) to study how adaptation alters the propagation and distribution of a single species in a linearly varying environment. Our goal is to better understand the propagation dynamics according to adaptation potential for a single species and to derive useful approximations for limit-adaptation cases. We address the question of how adaptation potential affects the geographical dynamics of a single species by providing approximations of species propagation speed under extreme scenarios (very low or very high adaptation potential) and using extensive simulations to understand intermediate scenarios. We link the local adaption and limits to range size to the variation in propagation speeds.

## 2. Kirkpatrick and Barton’s model for a single species’ range evolution along a linear gradient

The one-species model proposed by Kirkpatrick and Barton (1997) is a spatially explicit model in heterogeneous space accounting simultaneously for migration effects and adaptation. It assumes individuals are characterized by a phenotypic trait and that heterogeneity in space is given by a continuous cline of the optimal value for this phenotype. Individuals whose phenotype deviates from this optimum will suffer a fitness penalty. Although this model assumes that the environmental cline remains fixed in time, it provides a framework to study, for example, the effects of climate change on species distributions (see e.g. Norberg et al. (2012)), as it can easily be modified to include a time-varying environment. It is also suitable to study invasion scenarios, linking the characteristics of the environment and those of the introduced population.

The model assumes infinite one-dimensional linear space and considers local population density *n*(*t, x*), i.e. the density of individuals at location *x* at time *t* ≥ 0, and the mean phenotypic value of this population, 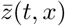, at this time and this location. The environmental cline is modeled through the optimal phenotype function *θ*(*x*) = *Bx* meaning the optimal value varies linearly through space. After a renormalization of the original variables and parameters in the full system (see Kirkpatrick and Barton (1997) for details), the equations governing density and phenotype dynamics are given by

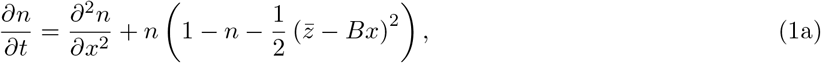

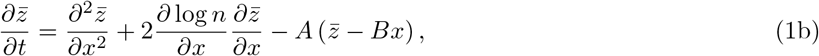

where *A* is a measure of *adaptation potential* of the species (*A* is proportional to genetic variance) and *B* is the rate of change of the optimal phenotype through space, also considered to be a measure of *spatial heterogeneity*.

System (1) describes the eco-evolutionary dynamics of the species under local adaptation and spatial diffusion. The first term of Equation (1a) models the dispersal of the population through a diffusion process. The second term contains the local ecological dynamics, corresponding to the logistic model and a penalizing term that captures local maladaptation. The first term of equation (1b) models the diffusion of genes that is linked to the diffusion of individuals, while the second term corrects for asymetries in gene flows (gene flow being more important from large populations to small populations than the other way round). The third term corresponds to the effects of local adaptation due to directional selection, driving the mean phenotype value 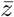 toward the local optimum *Bx* at a rate *A*.

Migration and adaptation potential *A* have antagonistic effects, whose results vary depending on the spatial heterogeneity *B*. Depending on *A* and *B*, the population may survive in a limited space (for intermediate values of *A* and *B*), may invade the whole space (when adaptation is larger than a certain critical value, allowing the population to surmount spatial heterogeneity) or may become extinct (when adaptation is too small with respect to spatial heterogeneity; (Kirkpatrick and Barton (1997))). This result can be partially re-stated in terms of propagation speeds (Fisher (1937)), which answer at the same time the question of geographical dynamics of the population: if we consider as initial condition a geographical frontier, i.e., the initial condition is *n*(0, *x*) = 1 for *x* ≤ 0 and 0 otherwise, with the species being perfectly locally adapted 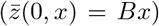 wherever it is present (*n*(0, *x*) = 1), then the solutions behave like propagating fronts with a characteristic speed. For Kirkpatrick and Barton’s one-species model, the direction and magnitude of the advancing front depend on the parameters *A* and *B*. Positive speeds mean the front moves towards positive values of *x* so that the species progressively invade (hereafter *invasion fronts*). On the contrary, negative values mean that the species distribution retracts (either to a limited range or toward the extinction of the species, hereafter *extinction fronts*). We dub *c*_*KB*_ (*A, B*) the speed of the solution of system (1) for parameters *A* and *B*.

In terms of propagation speeds, species whose borders correspond to invasion fronts are able to continuously adapt to new environments and thus will always be able to invade the whole space. On the contrary, negative speed fronts only mean maladapted gene flow is stronger than adaptation, causing local extinctions that can lead to two outcomes: either the population becomes extinct, or two fronts from different directions collide canceling out maladaptations in the center and allowing the species to survive in a limited space. We cannot distinguish between these two last outcomes based on speed alone, another demographic criterion is needed to do so. Refer to Figure 1 for a clearer link between propagation speeds and spatial distribution.

## 3. Explicit approximation of propagation speeds under various adaptation scenarios

We investigate the variation of propagation speeds for different values of parameters *A* and *B*, with a focus on the two limit cases of infinitely strong adaptation and very weak adaptation potentials. Even though it is unlikely species adapt infinitely fast, the variation in propagation speeds between these two limit cases can tell us when a finite adaptation is strong enough so that it is qualitatively infinite.

One first important result is that when adaptation goes to infinity, *A* → ∞, the system (1) becomes the Fisher-KPP equation (after Fisher (1937) and Kolmogorov et al. (1937), see the Appendix A.1 for details), given by

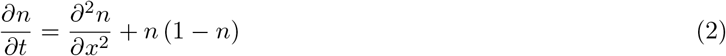

in its non-dimensional form (refer to the appendix for details on this infinite adaptation limit). Its solutions are traveling fronts with a minimal admissible speed of *c*_*F*_ = 2 (or, in its dimensional form, 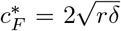 with *r* corresponding to the intrinsic growth rate of the population and *δ* a measure of its dispersal), so that for infinite adaptation potential invasion speed is finite and constant. Equation (2) has an infinity of solutions for different front speeds *c* ≥ *c*_*F*_, but *c*_*F*_ is the smallest one and the only one with biological meaning.

We can draw two other important conclusions thanks to equation (2). First, in an ecological context, the Fisher-KPP equation can only model propagation of species whose adaptation is so fast that they are continuously well-adapted everywhere, since the equation is the same as the system (1) neglecting maladaptation (and all the terms involving the phenotypic trait). Second, invasion speeds for the one-species model given by system (1) will always be lower than *c*_*F*_ = 2, since growth rate in the Fisher-KPP model is always larger than the one of the KB model, because maladaptation effects can only decrease population fitness (having thus a negative effect on speed). This means that the maximum speed of range expansion is only constrained by the species growth rate and dispersal ability (since it is 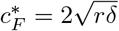 in its dimensional form).

At the other extreme of the adaptation gradient, the limit of small adaptations *A* → 0 needs to be studied more carefully. For *A* = 0, we would have a non-adapting species which cannot invade environments it is not suited to. We consider the term 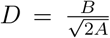 which we dub the *evolutionary challenge*, since it embodies the spatial heterogeneity to overcome for a given adaptation potential (measured not directly as *A*, but as 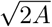). We consider species with decreasing adaptation potentials while keeping a constant evolutionary challenge (i.e. *A* → 0 with *D* constant). This provides a way to study a small adaptation potential while scaling the environment accordingly. This small adaptation limit has already been studied by Mirrahimi and Raoul (2013) and there is an explicit expression for the propagation speed for such low adaptation scenarios, given by:

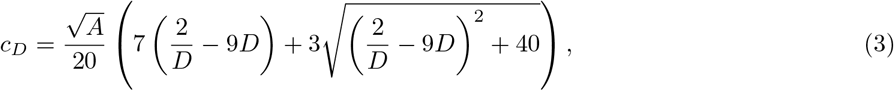

or 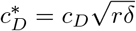 in its dimensional form.

Note that this expression is decreasing in *D*, meaning that for larger evolutionary challenges the invasion
speed will be smaller (refer to Figure 2). Also, invasion speed depends on the spatial heterogeneity (i.e., *B*) only through *D*. The value 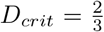 gives an invasion speed of 0, which means that for very small adaptation potentials, when the challenge is larger than *D*_*crit*_ the species goes extinct while when the challenge is smaller they are able to invade at a speed 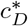. Notice how 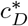 is proportional to 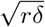, as is Fisher-KPP’s speed, corrected by a factor *c*_*D*_ that depends on the adaptation potential 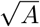 and challenge *D*, which corresponds to a loss in invasion efficiency due to maladaptation effects.

## 4. Relating propagation speeds to adaptation regimes

A natural question is then how these extreme scenarios relate to Kirkpatrick and Barton’s one-species model in terms of propagation speeds, which provides a method to concretely determine what strong and weak adaptations mean. To understand this, we numerically approximate the solution of the one-species model for a variety of parameter pairs (*A* and *B*) and compare the results to those given by the extreme adaptation limits (Figure 3. Refer to the Appendix A.2 for details on the numerical scheme).

**Figure 1:**
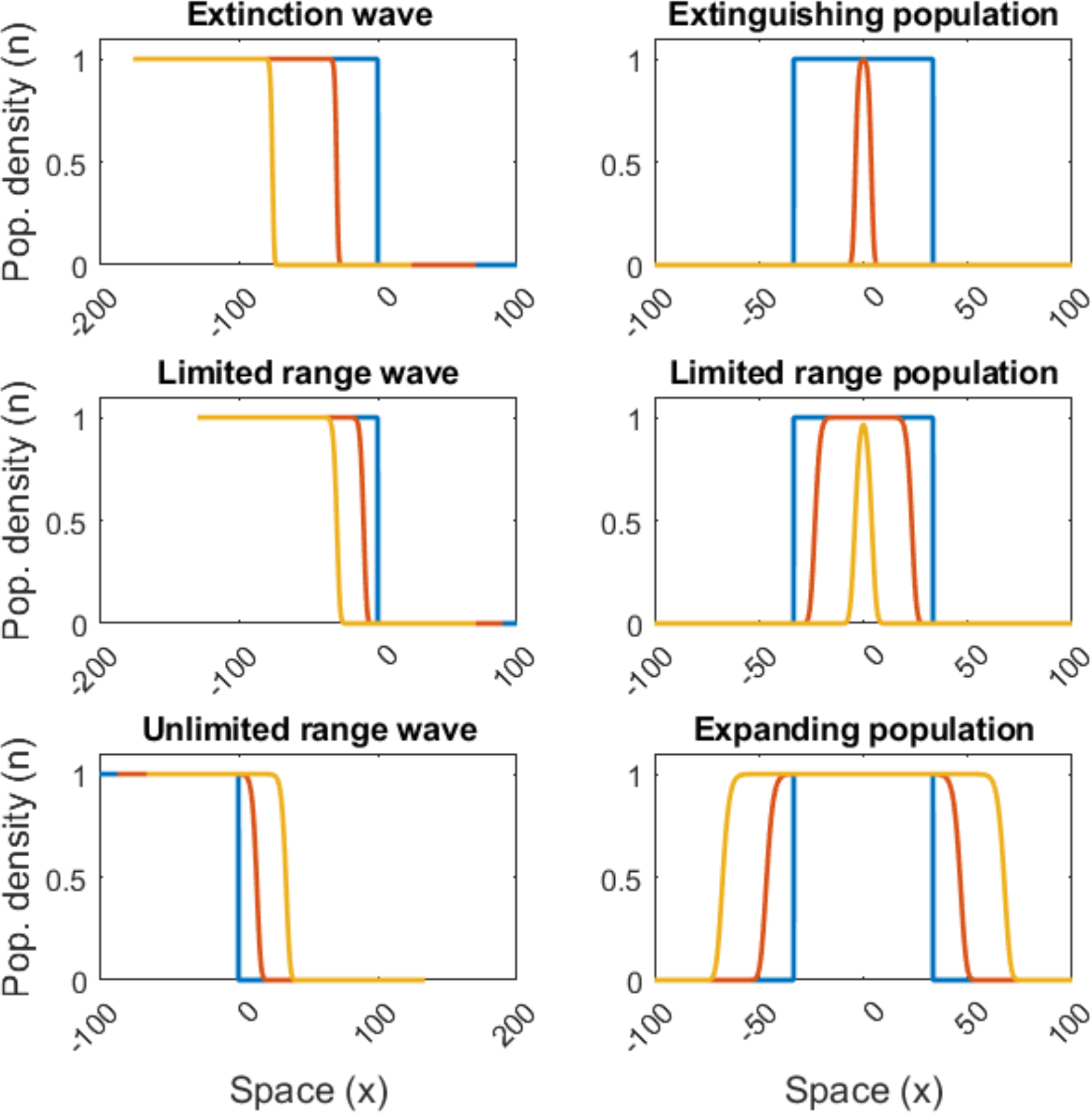
Panels showing the relation between a propagation wave and the respective population density distribution. In every panel, color blue indicates the initial condition, color red indicates an intermediate value (*t* = 20) and color yellow a long time (*t* = 50) distribution. The panels on the left column feature the dynamics of a boundary, whereas the panels on the right column feature the dynamic of an initially limited-range population distribution, with the same parameters (*A*, *B*) for each row. The first two rows show that a negative propagation speed may drive a population towards extinction (first row) or to a limited range distribution (second row). The third row shows that a positive propagation speed leads to an unlimited range distribution.

**Figure 2:**
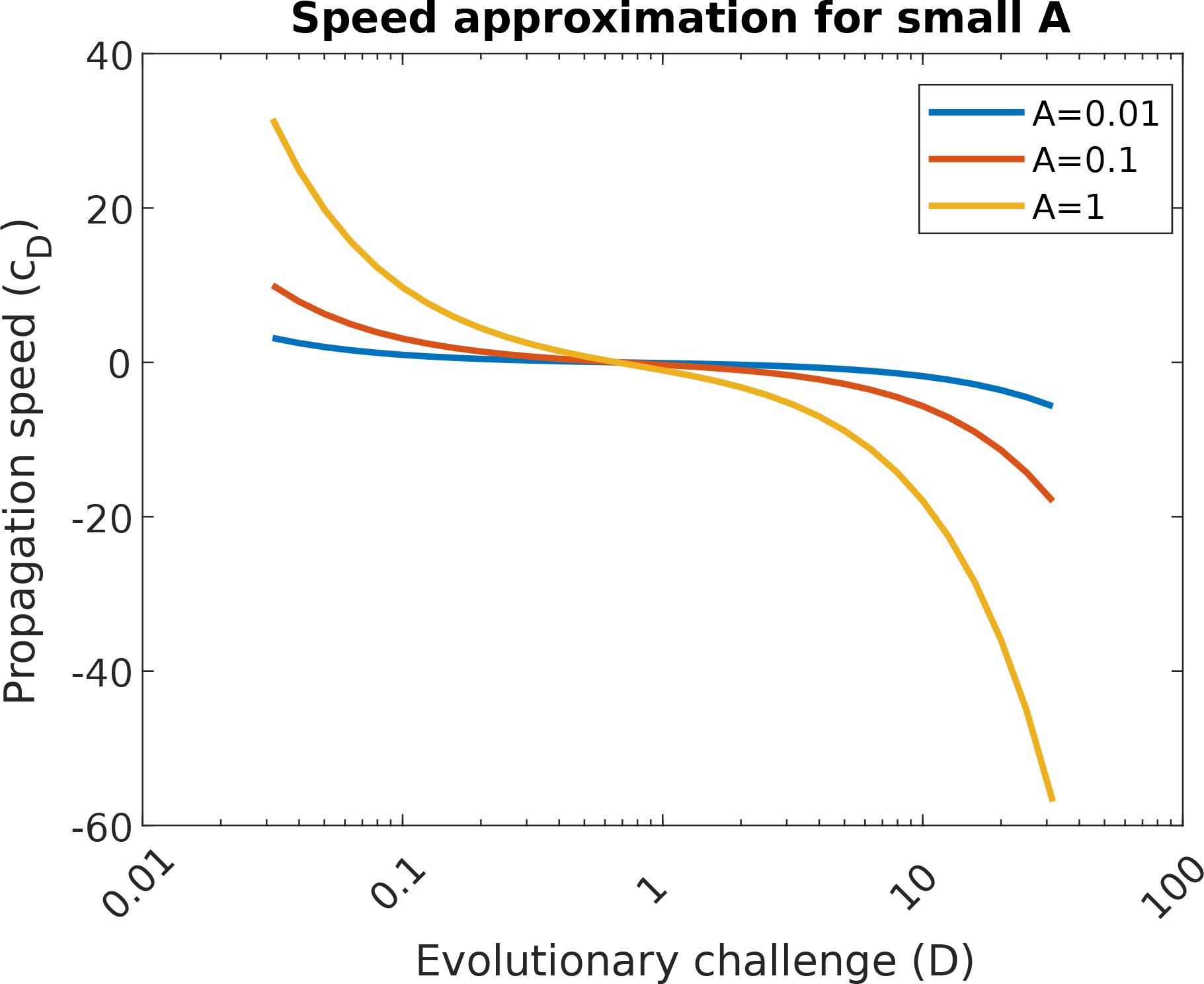
Propagation speed *c*_*D*_ as a function of the environmental challenge *D*, as defined by formula (3).

**Figure 3:**
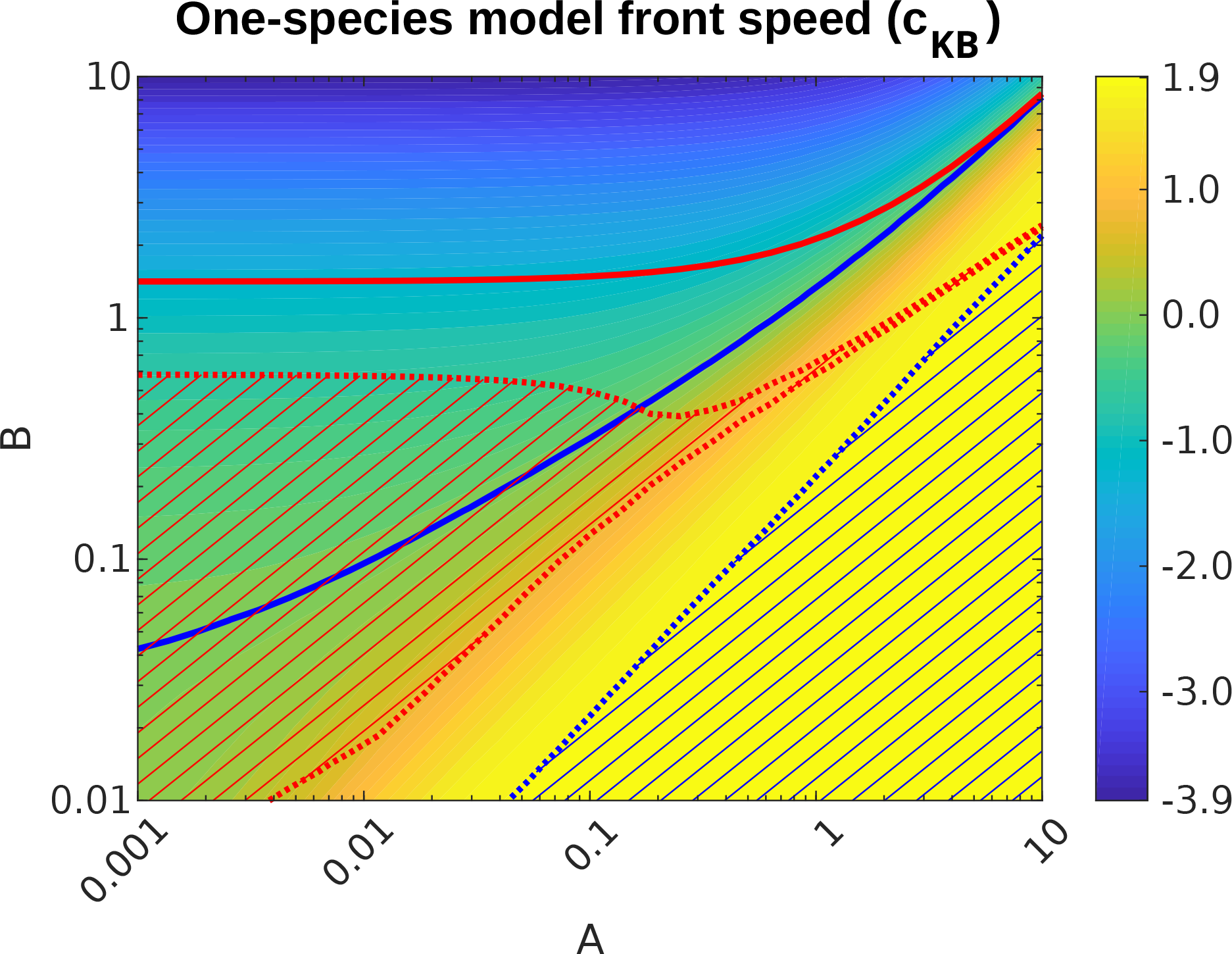
Approximated speeds for Kirkpatrick and Barton’s one-species model along with several important lines showing some regime changes in the original system (1). The color gradient shows the speed value for different (*A*, *B*) parameters (see color legend on the right-hand side of the plot). The thick red line shows the regime change between extinction and limited ranges, as approximated by Kirkpatrick and Barton (1997). The thick blue line corresponds to the regime change between limited and unlimited range, which is also the zero-level line for the invasion speed (also in Kirkpatrick and Barton (1997), Figure 2). The blue hatched area is the zone in the parameter space where the difference between propagation speed in Kirkpatrick and Barton’s model and Fisher-KPP’s model is at most 0.1, marking the *strong adaptation* regime; the red hatched area is where propagation speed in Kirkpatrick and Barton’s model is well approximated (i.e. the difference is at most 0.1) by the formula by Mirrahimi and Raoul (2013), i.e., given by (3), marking the *weak adaptation* regime.

First, note that propagation speed is increasing as a function of *A* and decreasing as a function of *B*, which is intuitive since larger adaptation potential and smaller spatial heterogeneity imply species will invade more easily.

The blue hatched area in Figure 3 shows where the difference between the one-species model speed *c*_*KB*_ and Fisher-KPP’s model speed *c*_*F*_ is at most 0.1. In this sense we can say that the blue dotted line marks *the limit between strong and intermediate adaptation potential*. A simple linear regression lets us approximate this region analytically by the inequality *A* ≥ 10^0.65^*B*. In other words, whenever adaptation potential surpasses the critical value *A*_crit_ = 10^0.65^*B* maladaptation effects are negligible, and species invade at a maximal speed, well approximated by the Fisher-KPP model.

The red hatched area in figure 3 marks where the speed in the one-species model is close to the small adaptation limit speed given by (3) (i.e. the difference between the propagation speed and the speed given by this formula is at most 0.1), so that in this zone adaptation is weak; thus the red dotted lines establish *the limit between weak and intermediate adaptation potential*.

The thick blue line corresponds to the zero-speed line, marking the division between positive and negative speeds. In other words, this line corresponds to the limit between unlimited and limited range which was studied in Kirkpatrick and Barton (1997).

Interestingly, this leaves only a small zone of parameter space that cannot be explicitly approximated. Propagation speeds then need to be assessed numerically since we do not have explicit formulae for them. This parameter space corresponds to the area outside the extinction regime (below the thick red line) that is not hatched, which we dub the *intermediate adaptation potential* zone. We also decided not to consider the speeds in the extinction zone since behavior in this zone is not very interesting, although we were able to measure corresponding extinction speeds.

We can find an explicit approximation for this line (by fitting a two-degree polynomial on the level-line, using MATLAB’s methods) which is given by:

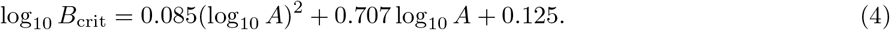

This means that for a given level of adaptation *A*, the population can only invade the whole space if the heterogeneity is smaller than *B*_crit_, otherwise it will suffer local extinctions, being restricted to a limited range or disappearing altogether. Solving the equation for *A*, we can interpret this result the other way round: for a given level of spatial heterogeneity *B*, the population will be able to invade only if its adaptation level is greater than the solution *A*\*. Unfortunately, the method we used does not provide us with an explanation or an intuition as to why the regime change occurs on this line, but it is nevertheless an improvement of the condition found by Kirkpatrick and Barton (1997), 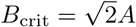, or equivalently,

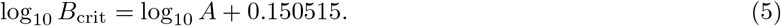

## 5. Discussion

Our model highlights how species adaptation can affect species extinctions and their geographical distributions. In this single-species model, explicit propagation speed and conditions of extinction can be obtained for most of the parameter space. These approximations highlight how different mechanisms act when considering low-vs high-adaptation potential.

The single-species adaptation model by Kirkpatrick and Barton (1997) shows various interesting behaviors. Even though there are no known explicit solutions to this system of equations, we were able to relate this model to other works, thereby providing explicit propagation speeds for most of the parameter space (refer to Figure 3 and its legend for details). Although the purpose of Kirkpatrick and Barton’s one-species model was not to study invasion processes, the usual approach to understand similar models is through the analysis of the speed of propagating fronts (as done in the first articles Fisher (1937) and Kolmogorov et al. (1937) or in literature in general, Skellam (1991), Shigesada and Kawasaki (1997)). This corresponds to the speed of ongoing local invasions (positive speed fronts) or local extinctions (negative speed fronts), which is the approach we took here. The original study by Kirkpatrick and Barton (1997) focused on the antagonistic effects of gene flow in a heterogeneous environment to understand the conditions under which a population has a finite geographical range. In order to comment on their results, it is helpful to recall the definitions of their compound parameters: the adaptation potential *A* = *G*/(2*V*_*s*_*r*\*) is the additive genetic variance of the population *G* divided by the basic population growth rate *r*\* and the strength of stabilizing selection (a smaller value of *V*_*s*_ meaning stronger selection); and the spatial heterogeneity 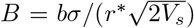 is proportional to the environmental gradient *b* and the dispersal rate *σ* (which makes sense since the more an individual disperses, the more the environment will be different proportionally to *b*). Although the approach developed by Kirkpatrick and Barton (1997) provided an efficient way to understand the determinants of range boundaries, it only deals with a small part of the parameter space (refer to Figure 3). Our results indicate that interesting conclusions, mostly about adaptation potential, can be obtained by looking at propagation speeds in the whole parameter range for *A* and *B*. Besides, the speeds of advancement or retraction of these models is interesting because it gives a way to roughly predict the future repartition of the modeled species (i.e. extinction/retraction or expansion of its range).

Our analysis revealed that adaptation potential, measured through parameter *A*, has a strictly positive effect on invasion speed. This speed is always smaller than that of Fisher’s model (Fisher (1937)), which neglects spatial heterogeneity. Thus, adaptation potential not only dictates whether a species can establish itself over space in a limited or unlimited manner, but it also helps overcome spatial heterogeneity, as shown by its invasion speed. For very strong adaptation potentials, the effects of spatial heterogeneity become negligible, with invasion speed being nearly equal to that of Fisher’s model. This is what we called the “strong adaptation zone” in Figure 3. For small adaptation potentials, the explicit approximation suggests that invasion speed critically depends on the evolutionary challenge 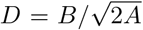 (equation (3)). This means that the fate of species with small adaptation
potential depends not so much on their adaptation capabilities, but rather on the spatial gradient to overcome given their evolutionary potential. The more challenging an environment is, the slower the invasion speed will be. Some case studies suggest that such constraints do act in nature. Consider for instance the reinvasion of its historic range by the California sea otter (*Enhydra lutris*) and the invasion of the sugar cane toad (*Rhinella marina*) in Australia. In the first case, Lubina and Levin (1988) showed important differences in expansion speeds at the north and south limits of the otters’ ranges possibly due to important environmental differences. In the second case, Urban et al. (2008) show that the invasion speed for the cane toad is not constant in time, and that periods of acceleration or decrease may be linked to changes in the environmental clines being invaded. In both cases, it is safe to suppose that the adaptation potential of the species is small, since both invasions started with only a few individuals.

Excluding the zone in Figure 3 where the species becomes extinct, the two approximations we used provide good descriptions of speeds in most of the parameter space. Having covered the infinite-and small-adaptation limits, only speeds in intermediate regimes remain to be studied. The explicit expressions for speeds we found make our model highly applicable. These explicit propagation speeds are also interesting because they let us determine where the limit between limited and unlimited range occurs. This limit matches the 0-speed isocline, corresponding to the zones where either increasing spatial heterogeneity or decreasing adaptation potential lead to species extinction. Limited ranges occur when two extinction fronts from opposite directions meet and maladaptation manages to cancel out in the middle, which is possible before population decreases critically if selection is not too strong. This is another motivation to study the intermediate-adaptation regimes more in depth.

We can draw two other important conclusions from this analysis: knowing a species adaptation potential *A* and the rate of change of its optimal phenotype over space, *B*, we can determine whether the species is going to invade space or not and at what speed. As a corollary, knowing the speed of advancement of a species and estimating the degree of spatial heterogeneity *B* can give an indirect assessment of the species adaptation potential *A*, which is directly related to its genetic variance. In other words, invasion speed can be used as an eco-evolutionary index allowing us to draw conclusions on genetic characteristics of a population. For instance, the previously cited cases of the sea otter and the cane toad (Lubina and Levin (1988) and Urban et al. (2008), respectively) are ideal cases to which our framework could be applied, for example to determine whether adaptation potential is the same along the different environments, which would explain the difference in invasion speeds only as a consequence of changing environments (i.e., different spatial heterogeneities *B*) and not due to genetic characteristics of the species in question.

The idea of using ecological emergent properties of a system to approximate evolutionary quantities echoes some approaches from evolutionary demography. In Hiltunen et al. (2014), prey evolution affects the phase diagram of consumer-resource oscillations. The authors propose, based on cycle observations alone, to compute an Evolutionary Dynamics Index quantifying ongoing prey evolution. In our case, having sufficient knowledge of the slope of the optimal niche (*B*) and of adaptation potential (*A*) lets us draw predictions on spatial dynamics. In Yoshida et al. (2003) it is also possible to infer characteristics of the population genetics based on the nature of the observed predator-prey cycles.

It would be valuable to use the explicit formula we provide to compare invasion speeds with those observed in nature. Such a work has already been done for numerical approximations of Kirkpatrick and Barton’s one-species model. García-Ramos and Rodríguez (2002) explored evolutionary speeds given by this model and compared them to the observed speeds for the expansion of the muskrat (*Ondatra zibethicus*) in Europe. They found empirical expansion speeds to be within the range predicted by the model. However, discrepancies have also been shown between observed expansion speeds and those predicted by the Fisher model (equation (2)), as remarked in Grosholz (1996), with speeds being either under-or overestimated. While these discrepancies may be due to an incorrect estimation of ecological parameters, we suggest other possibilities, such as limits due to lagging species adaptation or variation in species interactions.

In order to focus on the role of adaptation, we took a simple approach to ecological dynamics, relying on a simple logistic growth. In the context of species invasion, however, densities are low at the front, so that Allee effects may be commonly encountered. Petrovskii et al. (2002, 2005) showed that for a population model with Allee effects, it is not always possible to observe traveling waves and that various modes of propagation and persistence may be found (for example, patchy invasion). Burton et al. (2010) and Béenichou et al. (2012) also show that expanding fronts usually select for dispersive traits, so that invasion speeds are usually larger than predicted by constant diffusion models.

Changes in species distributions are nowadays commonplace, as species track changes in their environment and due to the accumulation of invasive species transported by human activities. Our results highlight how geographical shifts may rely on different mechanisms when species adaptation happens slowly or fast. Understanding the future of diversity depends on the development of models of co-evolving ecological networks in heterogeneous space, and the gathering of empirical data documenting simultaneously changes in species trait and distribution.

## A. Appendices

## A.1. Infinite adaptation case

We can show that when adaptation potential is high, i.e. when *A* → ∞, then the solution of the KB equations (system (1)) converges to the solution of the Fisher-KPP equation (equation (2)). This may seem intuitive, since population density is penalized by the maladaptation term 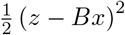, and when population adapts rapidly, this term should become negligible.

We propose that, for fixed values of the cline steepness *B*, when *A* → ∞ then the population density for the KB equations, *n*_*A,B*_, tends to the solution of the Fisher-KPP equations *n*_*F*_. We may write this as *n*_∞,*B*_ = *n*_*F*_. We propose the change of variables *w* = *z* − *Bx* so that equations (1a) and (1b) are rewritten as

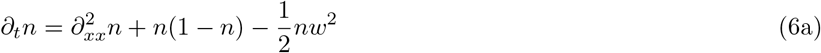

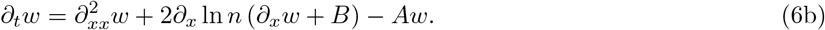

If we take equation (6b), multiply by *w* and integrate over *x*, we obtain (thanks to the integration by parts formula)

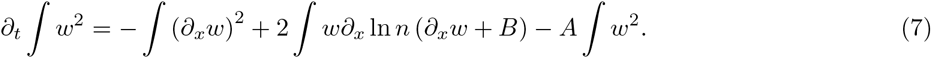

We wish to find estimates for the second term in the right hand side of this equation. We will assume that *∂*_*x*_ ln *n* is uniformly bounded over *A*, i.e., for fixed valued of *B*, |*∂*_*x*_ ln *n*(*t, x*)| ≤ *C* for every (*t*, *x*) with *C* a constant that does not depend on *A*.

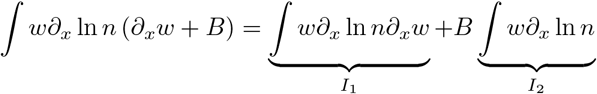

We can bound the first integral since

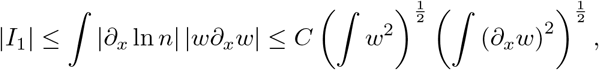

and thanks to Young’s inequality we can take some *ϵ* >0 so that

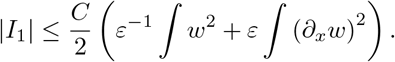

We need additional assumptions to obtain similar bounds on the integral *I*_2_. For example:

1. If additionally *∂*_*x*_ ln *n*(*t*, ·) ∈ *L*^2^(**R**) for every *A* and the *L*^2^ norms are uniformly bounded over *A*, say ∫(*∂*_*x*_ ln *n*)^2^ ≤ *C*_1_ and *C*_1_ does not depend on *A*,

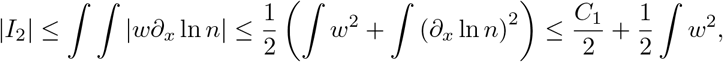

where we used Young’s inequality.
2. If additionally *∂*_*x*_ ln *n*(*t*, ·) ∈ *L*^1^(**R**) with a uniform bound over *A*, then we can use Hölder’s inequality in the following way

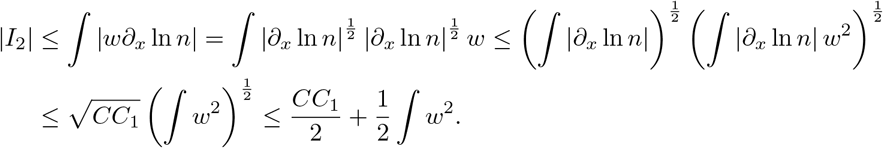
3. If we do not make additional assumptions on *∂*_*x*_ ln *n* but we suppose for example that we can control the *L*^1^ norm of *w* by its *L*^2^ norm, and the bound is uniform over *A*, we have

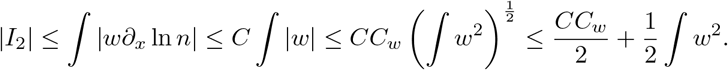

In any case, we found a bound of the form 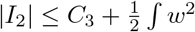, with *C*_3_ not depending on *A*.

Replacing the previously found bounds on expression (7) we find that

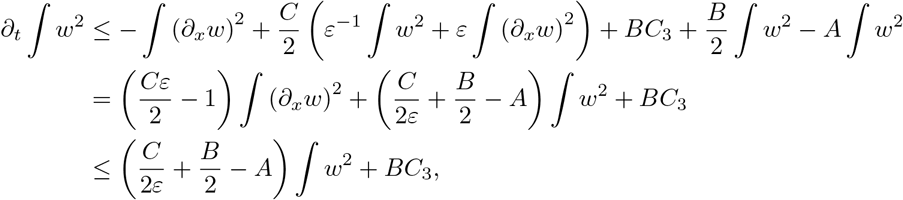

which is true for *ϵ* small enough (for example, 0 < *ϵ* ≤ *C*^−1^). Defining 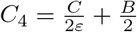, this implies that

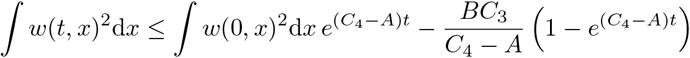

which tends to 0 for every *t* when *A* tends to *∞*. This implies in turn that 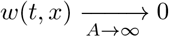.

We conclude that for any value of *B*, the function *w*(*t*, *x*) tends to 0 when *A* tends to infinity. This implies that, in the limit, the population density *n*_∞,*B*_ satisfies the Fisher-KPP equation, so *n*_∞,*B*_ = *n*_*F*_. In other words, the Fisher-KPP equation can be seen as a case of the KB equations when population adaptation potential is infinitely high.

## A.2. Numerical schemes

We present here the discretization we used to approximate the propagation speed in Kikpatrick and Barton’s model. Since it is already cumbersome to analyze it for the one-species model, for the two species model we only present the used scheme and describe briefly the problems we encountered.

As usual for a finite differences scheme, we consider a discretization of a finite time interval [0, *T*] and a time step ∆*t*, giving a time mesh *t* = 0, *t*_1_ = ∆*t*, etc., with the general formula *t*_*k*_ = *k*∆*t*, *k* ≥ 0; we also consider an one-dimensional space interval [−*L*, *L*] and a fixed spatial step ∆*x* so that we have mesh points *x*_*l*_ = −*L* + *l*∆*x*, *l* ≥ 0.

When considering an explicit time-forward scheme, we find the system

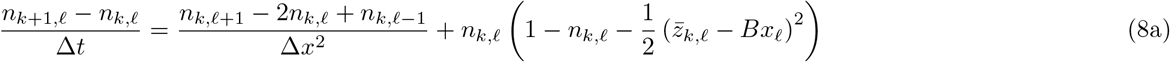

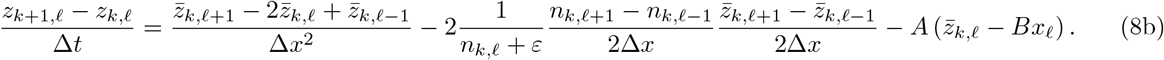

Notice that the solution for *n*_*k*+1_,*l* in terms of the *n*_*k*_ is almost a convex combination of these terms, explicitly

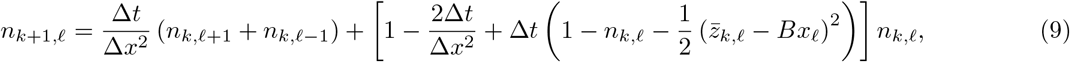

for it to be a (sub-)convex combination of the solution at different points of the mesh at the instant *t*_*k*_, we need the coefficients to be greater than zero and for their sum to be at most 1. Supposing that any desirable solution satisfies 0 ≤ *n*_*k,l*_ ≤ 1 for any *k*, *l* ≥ 0, the first condition is verified to be true if

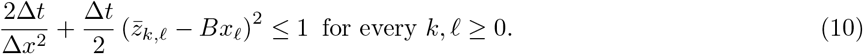

It is difficult to predict the values of 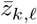 since it is ill-defined whenever *n* = 0 and simulations show instabilities when the *n*_*k,l*_ are close to zero and the mesh is not well chosen, however, the well working cases show that at the front tip there is an almost constant distance between 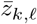 and the optimal phenotype *Bx*_*l*_. Since in some simulations we imposed 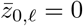 everywhere, and that locally this distance tends to decrease when 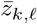 is too far from the optimum, then |*z*_*k,l*_ − *Bx*_*l*_ cannot be bigger than *BL* for a sufficiently big spatial window [−*L*, *L*]. We find thus that if the condition

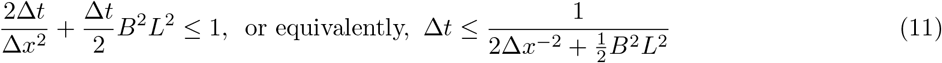

is met, then the stability condition (10) is valid.

Notice that when ∆*x*^−2^ is big enough compared to *B*^2^*L*^2^ then condition (11) is just the usual CFL-condition for the stability of explicit finite differences schemes for reaction-diffusion equations. However if *B* and *L* are bigger, this stability condition becomes highly restrictive. This actually made explicit schemes of this kind unpractical for our study.

We proposed our own non-linear implicit scheme for Kirkpatrick and Barton’s equations given, in the same presented mesh, by the following equations:

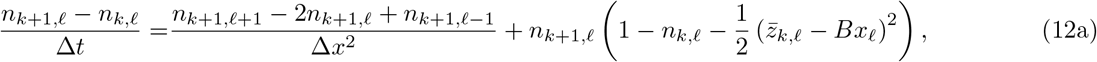

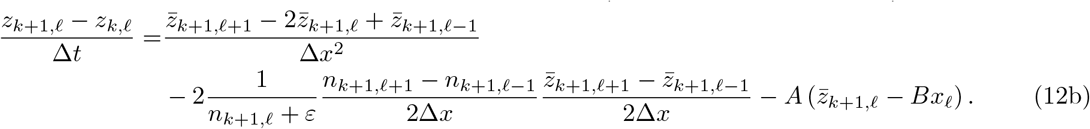

Notice that, for each time point *t*_*k*+1_, equation (12a) is linear on the vector *n*_*k*+1_ given that the values *n*_*k,l*_ and 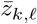 are known for each *l* ≥ 0 and thus it may be solved by matrix inversion techniques, with coefficients depending on the solution at previous time step *t*_*k*_. Once the vector *n*_*k*+1_ is known, the equation for the vector 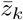 is just a linear one (with time-varying coefficients) that can also be solved with matrix inversion techniques.

Although we did not study the stability of the finite differences scheme (12), it behaved well for reasonable mesh parameters, and we were thus able to approximate the propagation speeds for a large family of (*A, B*) values.

